# ARABIDOPSIS NITRATE REGULATED 1 acts as a negative modulator of seed germination by activating *ABI3* expression

**DOI:** 10.1101/708222

**Authors:** Jia-Hui Lin, Lin-Hui Yu, Cheng-Bin Xiang

## Abstract

Seed germination is a crucial transition point in plant life and is tightly regulated by environmental conditions through the coordination of two phytohormones, gibberellin and abscisic acid (ABA). To avoid unfavorable conditions, plants have evolved safeguard mechanisms for seed germination. Here, we report a novel function of the Arabidopsis MADS-box transcription factor ARABIDOPSIS NITRATE REGULATED 1 (ANR1) in seed germination. *ANR1* knockout mutant is insensitive to ABA, salt, and osmotic stress during the seed germination and early seedling development stages, whereas *ANR1*-overexpressing lines are hypersensitive. ANR1 is responsive to ABA and abiotic stresses and upregulates the expression of *ABI3* to suppress seed germination. *ANR1* and *ABI3* have similar expression pattern during seed germination. Genetically, *ABI3* acts downstream of *ANR1*. Chromatin immunoprecipitation and yeast-one-hybrid assays showed that ANR1 could bind to the *ABI3* promoter to regulate its expression. In addition, ANR1 acts synergistically with AGL21 to suppress seed germination in response to ABA as evidenced by *anr1 agl21* double mutant. Taken together, our results demonstrate that the ANR1 plays an important role in regulating seed germination and early post-germination growth. ANR1 and AGL21 together constitutes a safeguard mechanism for seed germination to avoid unfavorable conditions.

## INTRODUTION

A successful transition of a seed into a seedling is a prerequisite for plant propagation and crop yield. Such transitions are mainly based on optimized and controlled molecular mechanisms that break seed dormancy and initiate the process of germination and seedling establishment (Rajjou *et al*., 2012; Miransari & Smith, 2014). Seed dormancy is crucial for it prevents seeds from competing among species, germinating under unfavorable conditions (Finkelstein *et al*., 2008; Shu *et al*., 2016). Seed germination is a term applied for multiple sequential mechanisms occurring inside a seed, starting from water imbibition, followed by embryo-activation, leading to elongation and expansion of embryo, and finally the emergence of radical out of the seed coat (Hermann *et al*., 2007). Extensive and intensive studies have been dedicated towards solving the mysteries of the duo of seed dormancy and germination.

The end of seed dormancy or initiation of germination is a vulnerable phase in a plant life, which is prone to suffer adverse environmental stresses (Lopez-Molina *et al*., 2002; Rajjou *et al*., 2012). The process of germination is primarily regulated by plant hormones like abscisic acid (ABA), gibberellic acid (GA), auxin, ethylene, and brassinosteroids (BR) (Finkelstein *et al*., 2002; Bai *et al*., 2012). The role of ABA is well-demonstrated in the regulation and maintenance of seed dormancy as well as the checkpoints for arresting seedling development (Lopez-Molina *et al*., 2001). Initially, ABA is accumulated in developing embryo which further regulates the seed dormancy, seed growth, and seed maturation (Finkelstein *et al*., 2002; Nambara & Marion-Poll, 2003). However, it rapidly diminishes in germination (Gubler *et al*.,2005). Plant hormones ethylene, gibberellins, brassinosteroids and cytokinins positively regulate seed germination (Kucera *et al*., 2005; Hermann *et al*., 2007), whereas ABA positively regulates the seed dormancy through a well-defined genetic signaling pathway (Finkelstein *et al*., 2008).

ABA Insensitive (*ABI*) genes are the core performers in the ABA-mediated signaling pathway in Arabidopsis. Group A of *ABI* genes (*ABI1* and *ABI2*) constitute type-2C protein phosphatases (PP2Cs) that are negative regulators in the ABA signaling pathway. Although the group B of *ABI* genes (*ABI3, ABI4* and *ABI5*)contributes in overlapping functions, they encode three distinct transcription factors, domain B3, AP2, and bZIP, respectively (Parcy *et al*., 1994; Finkelstein *et al*., 1998; Finkelstein & Lynch, 2000). Allelic mutants of group A (*abi1-1* and *abi2-1*)demonstrate reduction in dormancy (Koornneef *et al*., 1984; Ma *et al*., 2009), while group B mutants (*abi3, abi4* and *abi5*) show phenotypes insensitive to ABA during germination and early seedling development (Finkelstein *et al*., 1998; Finkelstein & Lynch, 2000; Soderman *et al*., 2000).

*ABI3* is responsive to ABA in early development of seeds (Giraudat *et al*., 1992; Parcy *et al*., 1997; Nambara *et al*., 2000; Zhang *et al*., 2002) and regulates a number of seed developmental genes that are primarily involved in thermo-inhibition and seed dormancy (Koornneef *et al*., 1984; Parcy *et al*., 1994; Tamura *et al*., 2006). At the transcriptional level, *ABI3* expression is promoted by LEAFY COTYLEDON1 (LEC1), LEC2, FUSCA3 (FUS3) and by ABI3 itself (To *et al*., 2006). However, the expression of *LEC1, LEC2* and *FUS3* terminates prior to the final embryo maturation, but the expression levels of *ABI3* remain relatively high in hydrated seeds until final maturation stages (Baumbusch *et al*., 2004; Park *et al*., 2011). During seed germination in response to ABA, *ABI3* is repressed by the chromatin modifier factor PICKLE (PKL) (Perruc *et al*., 2007). Furthermore, *ABI3* expression is strongly down regulated by DESPIERTO (DEP) (Barrero *et al*., 2010). WRKY41 also regulates *ABI3* expression (Ding *et al*., 2014). ABI3 directly enhances the expression of *ABI5* that subsequently interacted with *ABI3* and acts downstream of *ABI3* to execute an ABA-dependent growth arrest during germination (Lopez-Molina *et al*., 2002). A recent report shows that ABI5 forms a complex with ICE1-DELLA to fine tune ABA signaling in seed germination (Hu *et al*., 2019). ABI3 mediates dehydration stress recovery response in *Arabidopsis thaliana* by regulating expression of downstream seed-specific genes like *CRUCIFERIN1, CRUCIFERIN3* and *Late Embryogenesis Abundant* (*LEA*) proteins (Bedi *et al*., 2016). At the protein level, ABI3 is targeted for regulated proteolysis in the 26S proteasome by ABI3-INTERACTING PROTEIN 2 (AIP2) (Zhang *et al*., 2002). ABI3 is also targeted by PHYTOCHROME-INTERACTING FACTOR3-LIKE 5 (PIL5) transcription factor in imbibed seeds of Arabidopsis (Oh *et al*.,2009; Park *et al*., 2011). Due to the complexity of the molecular network modulating seed germination, there may be more unknown regulators of seed germination through regulating ABI3.

MCM1/AGAMOUS/DEFICIENS/SRF(MADS)-box transcription factors, with more than 100 members in plants (De Bodt *et al*., 2005), are the key regulatory factors involved in almost every development process of flowering plants (Smaczniake *et al*., 2012). However, their functions in seed germination and early post-germination growth are largely unexplored. Only few MADS-box genes were found involved in seed germination to date. *FLOWERING LOCUS C (FLC)/AGL25*, a key regulator of flowering, is involved in temperature-dependent seed germination by influencing ABA catabolic pathway and GA biosynthetic pathway (Chiang *et al*., 2009). *AGL67* functions as a potential negative regulator of seed germination, for *agl67* mutant accelerates the dissolution of seed dormancy (Bassel *et al*., 2011). Moreover, the MADS-box transcription factor *AGL21*, an important regulator of lateral root development (Yu *et al*., 2014), also acts as environmental surveillance of seed germination by regulating *ABI5* expression (Yu *et al*., 2017).

*ANR1/AGL44* was previously reported as a key determinant of nitrate induced lateral root development (Zhang & Forde, 1998). In this study, we show that *ANR1* is also expressed in seeds and regulates seed germination through ABA signaling pathway. Overexpression of *ANR1* conferred hypersensitive seed germination to ABA, high NaCl, and mannitol, while the *anr1* knockout mutants showed the opposite phenotypes under the same conditions. *ANR1* transcripts accumulated at high levels in dry seeds, while drastically reduced after cold stratification and imbibition, which is coincident with the expression pattern of *ABI3*. Biochemical and genetic analyses demonstrate that ANR1 regulates the expression of *ABI3* by directly binding to its promoter *in vivo*. Moreover, ANR1 functions synergistically with AGL21 to regulate seed germination as shown by the double mutant *anr1 agl21*. Our results show that ANR1 suppresses seed germination and post-germination growth by directly regulating *ABI3* in response to environmental stress signals as well as internal phytohormones. Therefore, ANR1 and AGL21 together afford seeds a double insurance to germinate only in favorable conditions.

## RESULTS

### *ANR1* expression pattern and response to multiple stresses

Previous preliminary experimental results showed that *ANR1* and *AGL21* had a similar expression patterns in root and embryos (Burgeff *et al*., 2002). To investigate the expression levels of *ANR1* in various organs, we carried out quantitative real-time PCR (qRT–PCR) and found that *ANR1* was highly expressed in roots and dry seeds, but with much lower expression levels in leaf and flower (Figure 1A). Subsequent expression analysis of *ANR1* in imbibed seeds at different germinating stages also revealed that *ANR1* transcripts accumulated at high levels in dry seeds, but substantially reduced upon the imbibition and drastically reduced to a very low level from 1 to 3 days after imbibition (Figure 1B), indicating a potential role of ANR1 in seed germination and early seedling development. The expression of *ANR1* increased at 7 and 10 days after seed imbibition probably because of root development of the seedlings (Figure 1B). As seed germination is strictly controlled by the antagonistic effect between ABA and GA (Yano *et al*., 2009; Yaish *et al*., 2010). We then examined the respond of *ANR1* to ABA and GA. qRT-PCR analyses showed that the *ANR1* expression level was significantly induced by exogenous ABA in wild type seeds at 1-3 days after imbibition (Figure 1C). The induction by ABA was fairly fast and significant upregulation was seen 1 hour after the treatment (Figure 1D). Moreover, we found that *ANR1* was also significantly upregulated by phytohormones IAA and MeJA as well as stress conditions (salt, osmotic stress, and nitrogen deficiency) (Figure 1E). All these results suggest that ANR1 is a potential modulator of seed germination responding to phytohormones (ABA, IAA and MeJA) and environmental signals.

**Figure 1.**
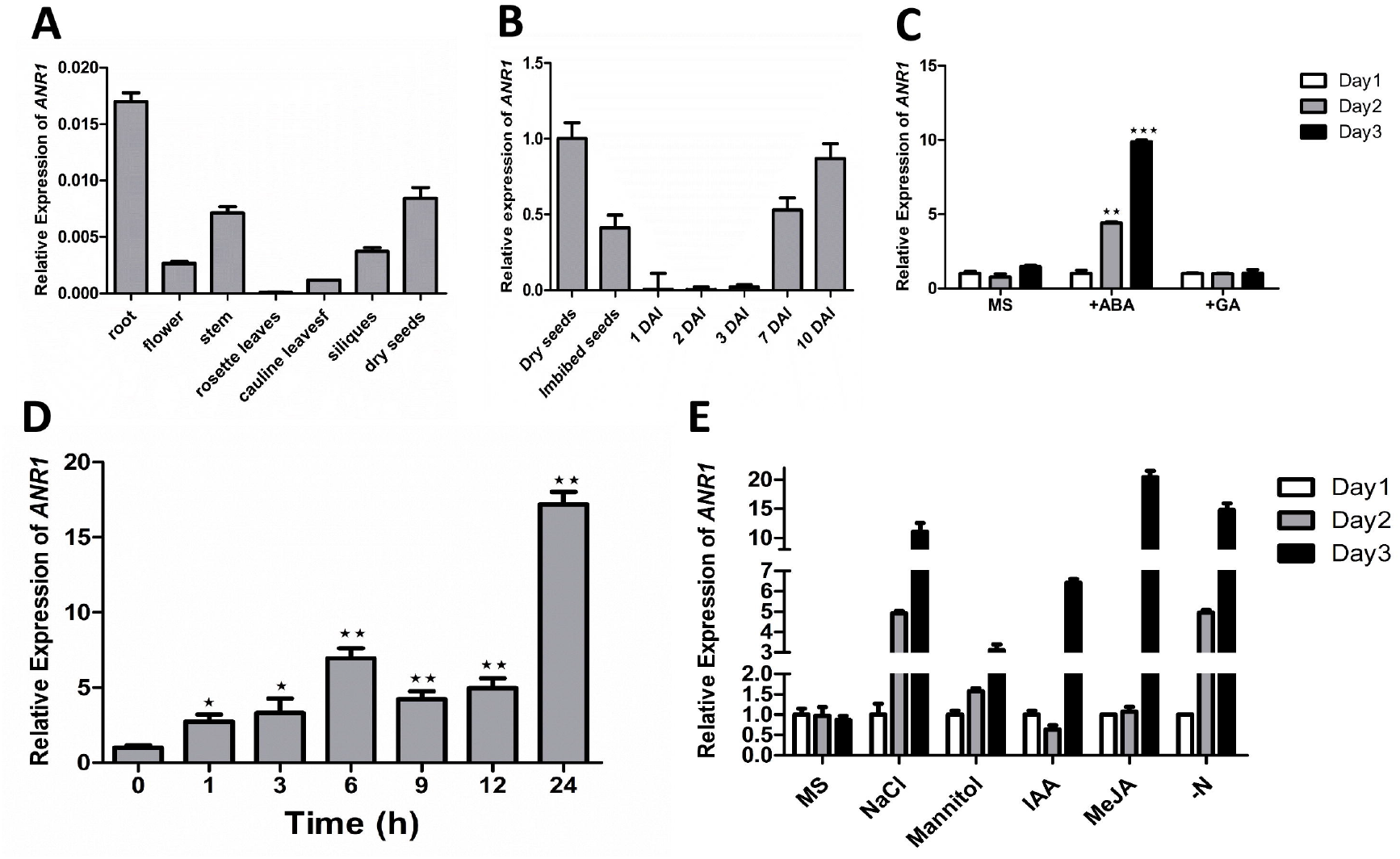
Expression of *ANR1* in seeds and seedlings. A. Expression of *ANR1* in different tissues of *Arabidopsis thaliana* wild type (Col-0 ecotype). *ANR1* transcript level was analyzed by qRT-PCR. Data are shown as mean ± SE (n = 3). B. Expression of *ANR1* in dry, imbibed, and germinating seeds. *ANR1* transcript level was analyzed by qRT-PCR in wild type plants (WT) during seed germination and early seedling development. WT weeds were imbibed and vernalized in water at 4°C for 48 h and grown on MS medium for different days (days after imbibition, DAI), and then the plants were harvested at the indicated time. Data are shown as mean ± SE (n = 3). C. Response of *ANR1* expression to ABA and GA in seeds. Vernalized wild type seeds were transferred to MS medium supplemented with 1 μM ABA or 5 μM GA and incubated for indicated time. *ANR1* transcript level was analyzed by qRT-PCR. Data are shown as mean ± SE (n = 3). *P < 0.05, **P < 0.01, ***P < 0.001. D. Time-course of ABA induction of *ANR1. ANR1* transcript level in 7-d-old WT seedlings treated with 20 μM ABA for different points was analyzed by qRT-PCR. Data are shown as mean ± SE (n = 3). *P < 0.05, **P < 0.01. E. Response of *ANR1* expression to various external signals. Vernalized wild-type seeds were transferred to MS medium supplemented with 100 mM NaCl, 250 mM mannitol, 10 μM IAA, 10 μM MeJA, and MS medium without nitrogen (-N) and then the plants were harvested at the indicated time. *ANR1* transcript level was analyzed by qRT-PCR. Data are shown as mean ± SE (n = 3). *P < 0.05, **P < 0.01.

### ANR1 acts as a negative modulator in seed germination and post-germination development responding to ABA, salt and osmotic stress

The *ANR1* expression pattern and its response to multiple environmental stress signals suggests that it has a potential physiological function in seed germination. To explore ANR1 function in seed germination, we generated *35S: ANR1* transgenic lines (OX4, OX10 and OX21) and obtained two T-DNA insertion mutants SALK_052716 (*anr1-1*) and SALK_043618 (*anr1-2*), which were confirmed by RT-PCR and qRT-PCR analysis (Supplemental Figure S1). Then we carried out germination assays using these lines. The germination rates of *35S:ANR1* and *anr1* were similar to that of the wild type under normal conditions (MS). However, when germinated on MS medium containing 1.0 μM ABA, *ANR1*-overexpressing lines were more sensitive to ABA, whereas the *anr1-1* and *anr1-2* mutants showed much higher resistance to ABA (Figure 2A, B and C). The cotyledon greening of *ANR1*-overexpressing lines and *anr1* mutants were indistinguishable from that of wild type in the absence of ABA, but hypersensitive and hyposensitive to ABA compared with wild type, respectively (Figure 2F). We further found that *ANR1*-overxpressing lines were more sensitive to salt and osmotic stress with lower germination rates and green cotyledon ratios. On the contrary, *anr1* mutants were much more resistant to the inhibition of salt and mannitol during seed germination and post-germination stages (Figure 2A, D, E and F). These data indicate that ANR1 acts as a negative regulator during seed germination and post-germination in response to ABA, salt and osmotic stresses.

**Figure 2.**
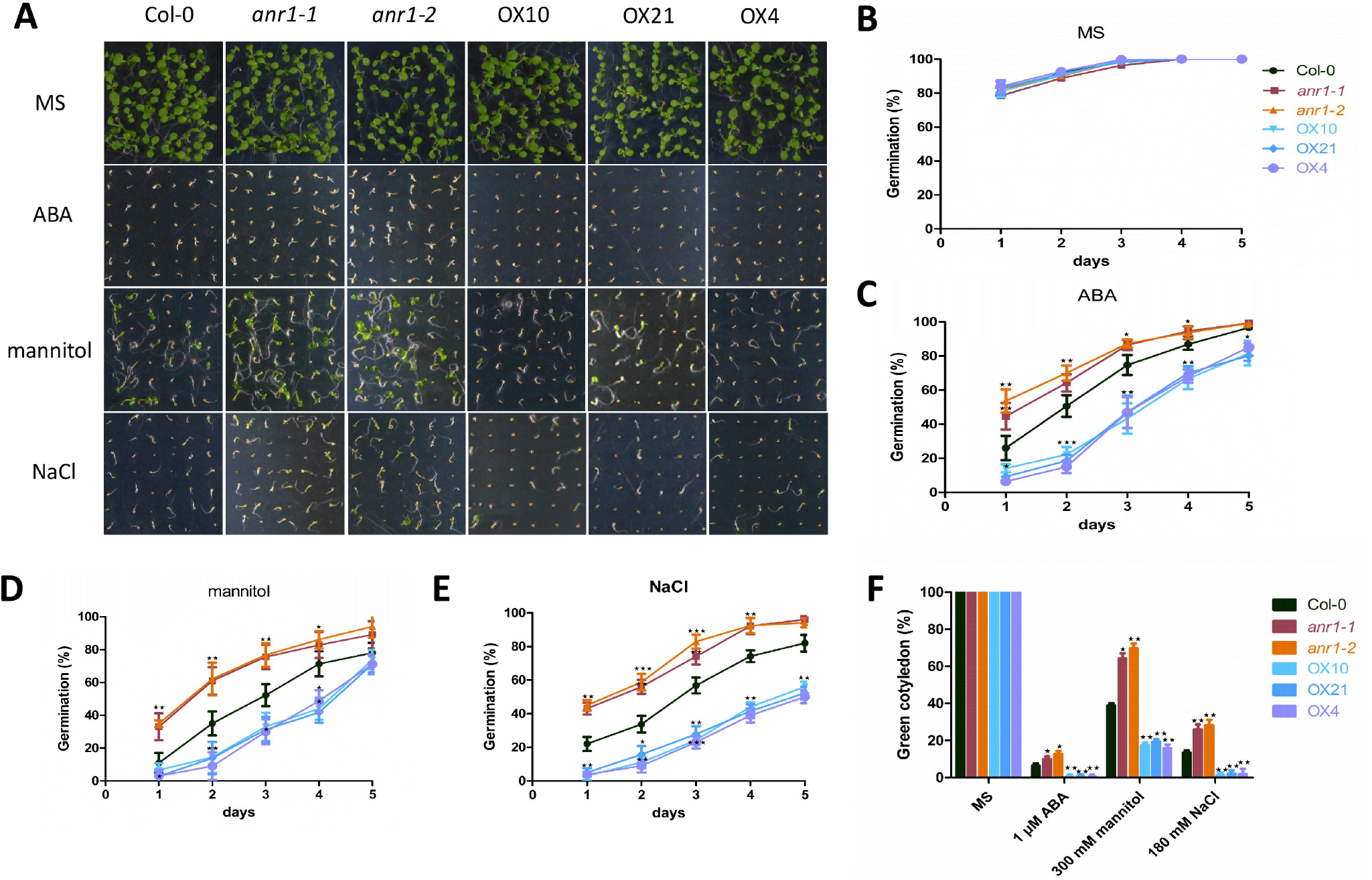
Response of *anr1* mutants and *ANR1*-overexpressing lines to ABA, mannitol and NaCl in seed germination. A. Seed germination phenotypes. Vernalized seeds of wild type (Col-0), knockout mutants (*anr1-1, anr1-2*), and *ANR1*-overexpressing lines (OX4, OX10, OX21) were transferred to MS medium or MS medium containing 1.0 μM ABA, 300 mM mannitol and 180 mM NaCl and grown for 8 days before the images were recorded. B-E. Seed germination curves. Seed germination rates were calculated at the indicated time for the lines and treatments in A. At least 42 seeds per genotype were measured in each replicate. Seeds from independent lines were used for replicates. Data are shown as mean ± SE (n = 3). F. Cotyledon greening ratio. Green cotyledon ratios were counted at day 12 after the end of vernalization. Data are shown as mean ± SE (n = 3). *P < 0.05, **P < 0.01.

### ANR1 is involved in ABA signaling

To study whether ANR1 plays a role in the ABA signaling pathway, we examined *ANR1* expression levels in *abi* mutants in response to ABA. We found that *ANR1* expression was significantly induced by exogenous ABA in the wild type, *abi3, abi4* and *abi5* mutant seeds. However, such induction was impaired in *abi1* and *abi2* mutant seeds (Figure 3), indicating *ANR1* may be act downstream of *ABI1, ABI2*, and upstream of *ABI3, ABI4, and ABI5* or parallel to them. To further investigated the role of ANR1 in ABA signaling, we analyzed the expression levels of several ABA-responsive genes in the *ANR1*-overexpressing lines and *anr1* mutants by qRT-PCR, including *ABI1, ABI2, ABI3, ABI4, ABI5, RD29A, RD29B, EARLY METHIONINE-LABELED 1 (AtEM1), SNF1-RELATED PROTEIN KINASE2.2* (*SnRK2.2*)and *SnRK2.3*. The results showed that ANR1 did not affect the expression of the upstream genes in ABA signaling pathway, such as *SnRK2.2, SnRK2.3, ABI1* and *ABI2*(Figure 4A-D). However, the downstream genes in ABA signaling pathway, such as *ABI3, ABI5* and *AtEM1*, were up regulated in *ANR1*-overexpressing seeds while significantly down regulated in *anr1* seeds (Figure 4E-4H). Another two ABA-inducible genes, *MYB DOMAIN PROTEIN 96* (*MYB96*) and *ACYL-COENZYME A-BINDING PROTEIN 1 (ACBP1*), which participating in the regulation of seed germination (Tran *et al*., 2007; Bu *et al*., 2009), were found having similar expression levels in *ANR1*-overexpressing and knockout seeds (Figure 4K and L). Taken together, ANR1 is likely involved in ABA signaling pathway, acting downstream of *ABI1, ABI2* and *SnRKs* while upstream of *ABI3*.

**Figure 3.**
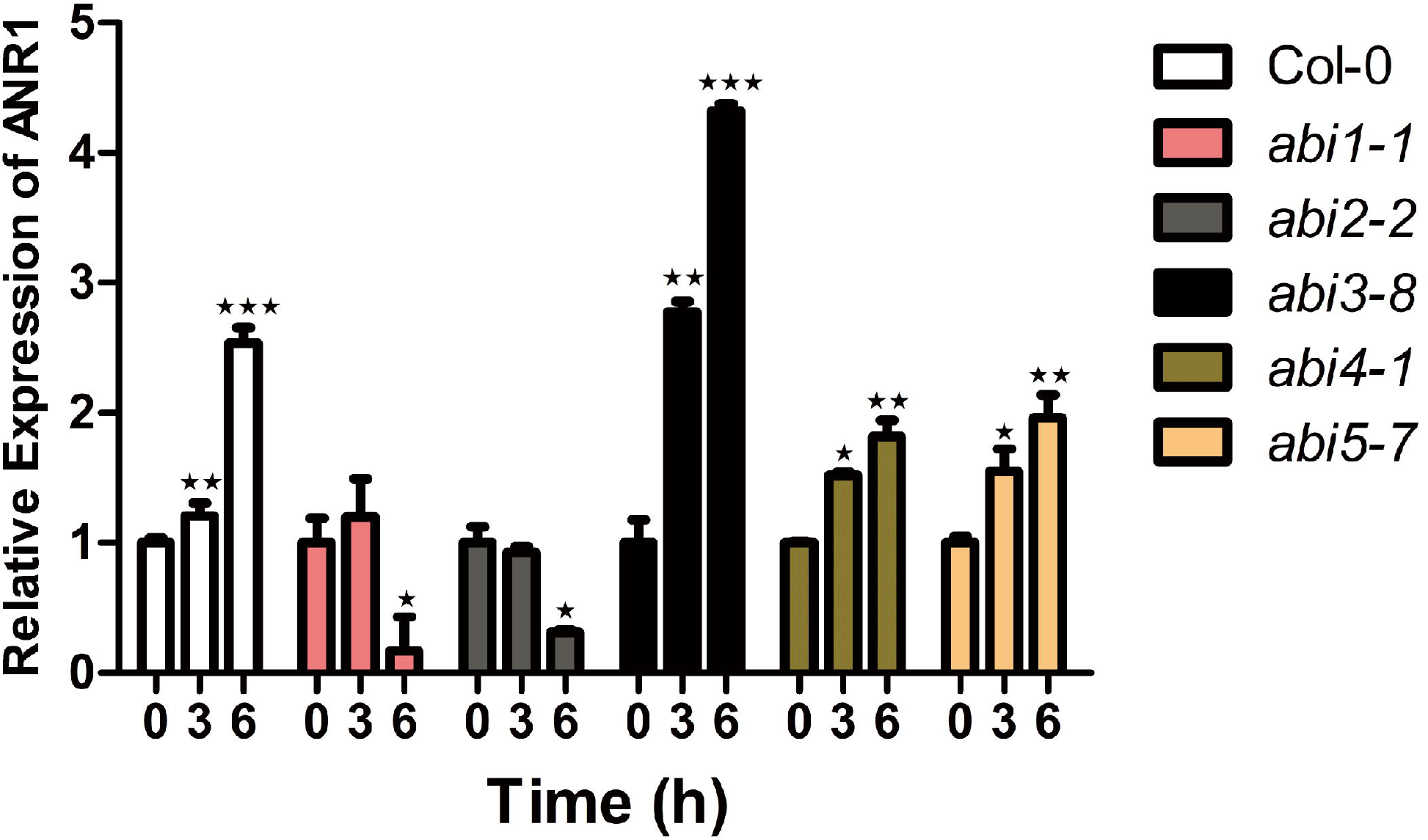
Expression of *ANR1* in WT and *abi* mutants in response to ABA. Response of *ANR1* expression to ABA treatments in WT and different *abi* mutants. Vernalized seeds of different genotypes were transferred to liquid MS medium supplemented with 1 μM ABA and then the plants were harvested at the indicated time for RNA extraction and real-time qPCR analyses. Data are shown as mean ± SE (n = 3). *P < 0.05, **P < 0.01, ***P < 0.001.

### ANR1 does not affect ABA and GA contents

To test whether ANR1 regulates seed germination by altering ABA or GA content in seeds, we first investigated the expression of ABA biosynthesis-related genes including *ABA DEFICIENT 1* (*ABA1*), *ABA2, ABA3, ABCISIC ALDEHYDE OXIDASE 3* (*AAO3*), *9-CIS-EPOXYCAROTENOID DIOXYGENASE* (*NCED3*) and ABA catabolism-related genes including *CYP707A1, CYP707A2, CYP707A3* and *CYP707A4*. The results showed that there were no significant expression differences of these genes between the wild type, *ANR1*-overepressing line, and *anr1-1* knockout mutant in the germinating seeds with or without ABA treatment (Supplemental Figure S2A-I). Consistent with these results, ABA content did not change in dry or imbibed seeds of *ANR1*-overepressing and knockout mutant compared with that in the wild type (Supplemental Figure S7A and B).

To test the sensitivity of seed germination to GA, we germinated the seed of wild type, overexpression lines (OX4, OX10 and OX21), and knockout mutants (*anr1-1* and *anr1-2*) on MS medium containing 0, 1 or 2 μM GA and found that there was no significant difference in germination among these lines (Supplemental Figure S3).

We also analyzed the expression levels of GA synthesis-related genes including *ENT - COPALYL DIPHOSPHATE* (*CPS*), *ENT KAURENE SYNTHASE* (*KS*), *ENT - KAURENOIC ACID OXIDASE 2* (*KAO2*), *GA20-OXIDASE* (*GA20OX1*), *GA20OX2, GA20OX3, GA REQUIRING 3* (*GA*), *GA3OX2* and *GA3OX3*, in the germinating seeds in presence or absence of ABA. No obvious difference was observed between the different genotypes as showed in Supplemental Figure S4. We further examined the expression levels of GA catabolism-related and GA signaling-related genes and found no significant difference among the genotypes (Supplemental Figure S5 and S6).

Consistent with these results, GA content did not change in dry or imbibed seeds of *ANR1*-overepressing and knockout mutant compared with that in the wild type (Supplemental Figure S7C and D).

### *ANR1* acts upstream of *ABI3* to regulate seed germination

Based on our previous data (Figure 4), we speculate that *ABI3* may be a target gene of ANR1. To confirm this, we compared the expression of *ANR1* and *ABI3* in seeds. As showed in Figure 5A, *ABI3* is highly expressed in dry seeds, and gradually decreased after 12–24 h of imbibition and then had a little increase after 48-72 h of imbibition. At 1–3 days on MS medium after imbibition, *ABI3* expression dropped to a very low level. Exogenous ABA obviously induced *ABI3* expression in germinating seeds (Figure 5A). It is noteworthy that *ABI3* was significantly up-regulated in *ANR1*-overexpressing while dramatically down-regulated in *anr1* seeds compared with wild type seeds (insets in Figure 5A), indicating *ABI3* expression is positively correlated with ANR1. We then checked the expression pattern of *ANR1*, and found that *ANR1* had a very similar expression pattern to *ABI3* in dry, imbibed and germinating seeds (Figure 5B). In addition, the expression pattern of *ANR1* and *ABI3* obtained from the eFP browser (Winter *et al*., 2007) also show that *ABI3* and *ANR1* has a very similar expression pattern in seeds after 0-72 h imbibition, with highest transcript level after 1h imbibition, then gradually declined after 3-24h imbibition (Figure 5C), consistent with our qRT-PCR data. Moreover, *ANR1* and *ABI3* showed alike expression pattern in developing seeds as showed in Figure 5D.

**Figure 4.**
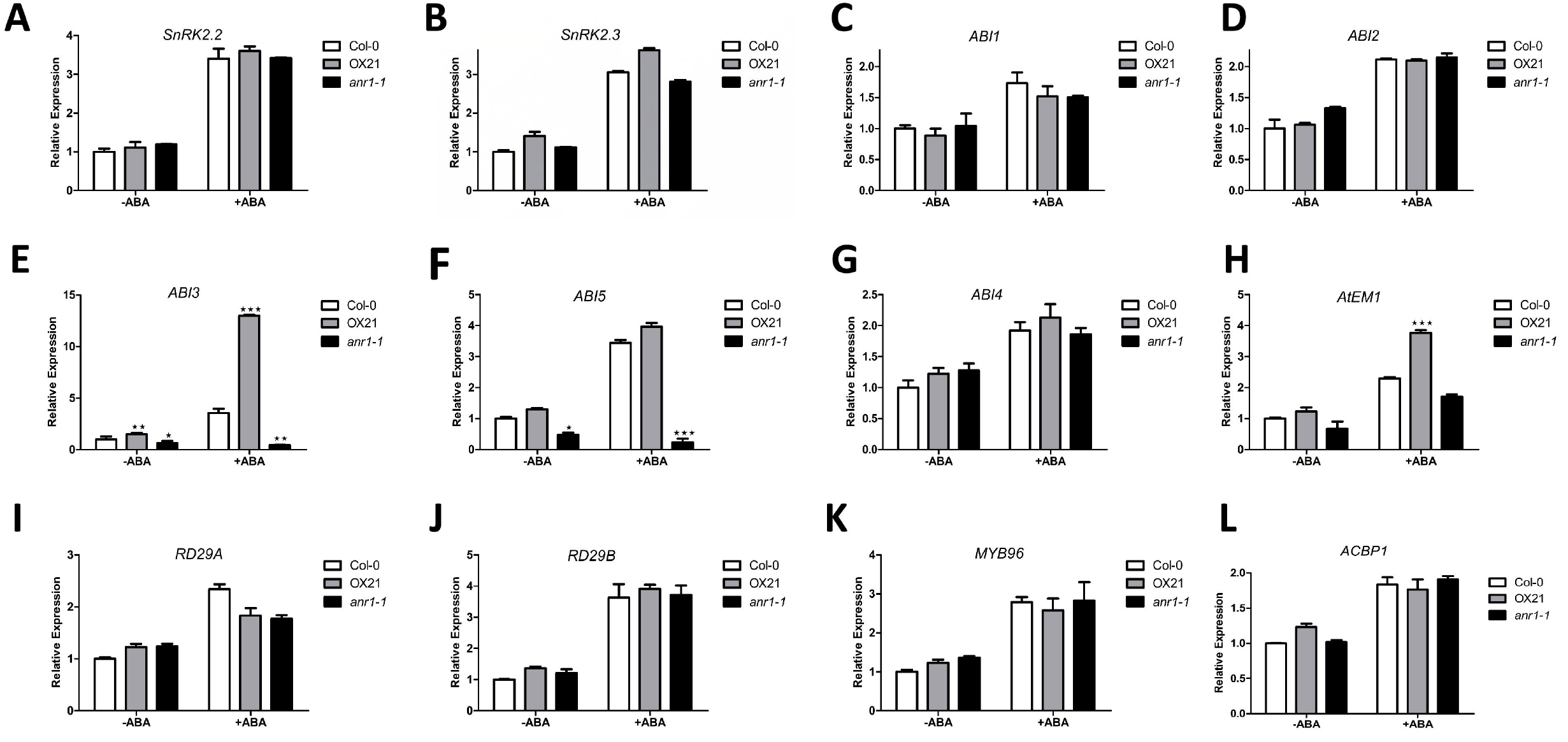
Expression of ABA-responsive genes in ABA-dependent seed germination of *anr1* mutants and *ANR1*-overexpressing lines. A-L. The vernalized seeds were germinated and grown on MS medium supplemented with or without 1 μM ABA and incubated for 3 days, and then the plants were harvested for RNA extraction. Transcript level of the gene indicated was analyzed by qRT-PCR. The *UBQ 5* gene was used as an internal control. Data are shown as mean ± SE (n = 3). *P < 0.05, **P <0.01.

**Figure 5.**
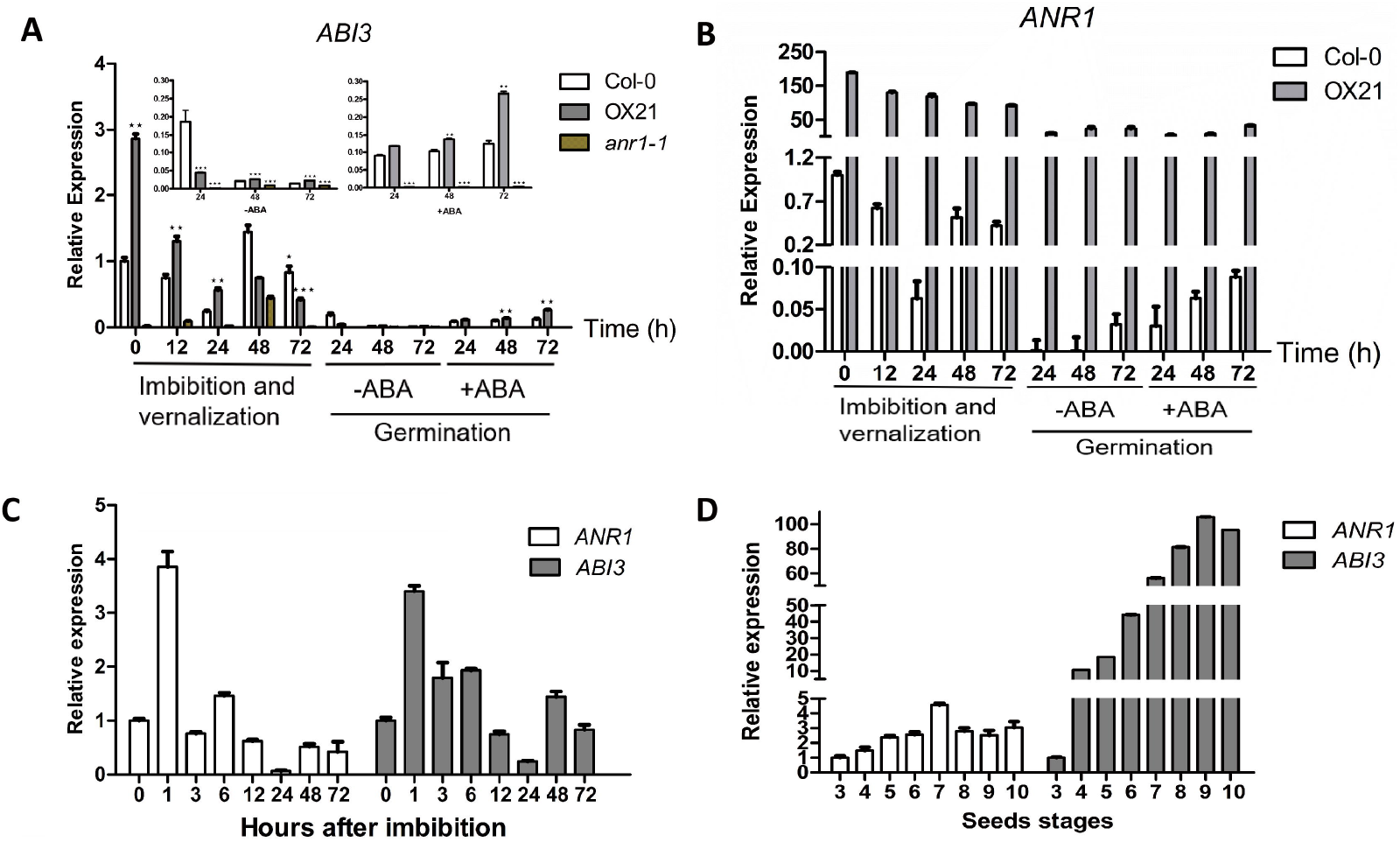
*ANR1* has similar expression pattern to *ABI3*. A. Expression of *ABI3* in wild type (Col-0), *ANR1*-overexpression line and *anr1* mutant. Seeds were imbibed and vernalized at 4°C for the indicated time before being sampled for RNA extraction. After 72 h imbibition, seeds were germinated on MS medium with or without 1 μM ABA for the indicated time before being sampled for RNA extraction. *ABI3* transcript levels were analyzed by qRT-PCR. The *UBQ5* gene was used as an internal control. Data are shown as mean ± SE (n = 3). *P < 0.05, **P <0.01, ***P < 0.001. B. Expression of *ANR1* in wild type (Col-0) and *ANR1*-overexpression line. Seeds were imbibed and vernalized in water at 4°C for different time. After 2 days of imbibition seeds were incubated on MS medium supplemented with or without 1 μM ABA, and then the seedlings were harvested at the indicated time for RNA extraction and real-time qPCR analyses of *ANR1* transcript level. Data are shown as mean ± SE (n = 3). C. The expression patterns of *ANR1* and *ABI3* in imbibed seeds obtained from the *Arabidopsis* eFP browser (Winter *et al*., 2007). D. Expression of *ANR1* and *ABI3* in wild type (Col-0). Seeds were imbibed and vernalized in water at 4°C for different times, and then the plants were harvested at the indicated time for RNA extraction and real-time qPCR analyses. Data are shown as mean ± SE (n = 3).

To confirm the relationship between *ANR1* and *ABI3*, we made crosses between the *ANR1*-overexpressing lines and *abi3-8* mutant and obtained the *ANR1-OX abi3-8* double mutant plants. Germination assay showed that all seeds had similar germination and green cotyledon ratios on MS medium. However, the *ANR1-OX abi3-8* double mutant was resistant to ABA, mannitol and NaCl in the seed germination and post-germination just like *abi3-8* mutant, while the *ANR1*-overexpressing seeds are hypersensitive under the same conditions (Figure 6). Taken together, our data indicate that *ANR1* is upstream of *ABI3* and *ABI3* is a potential target of ANR1.

**Figure 6.**
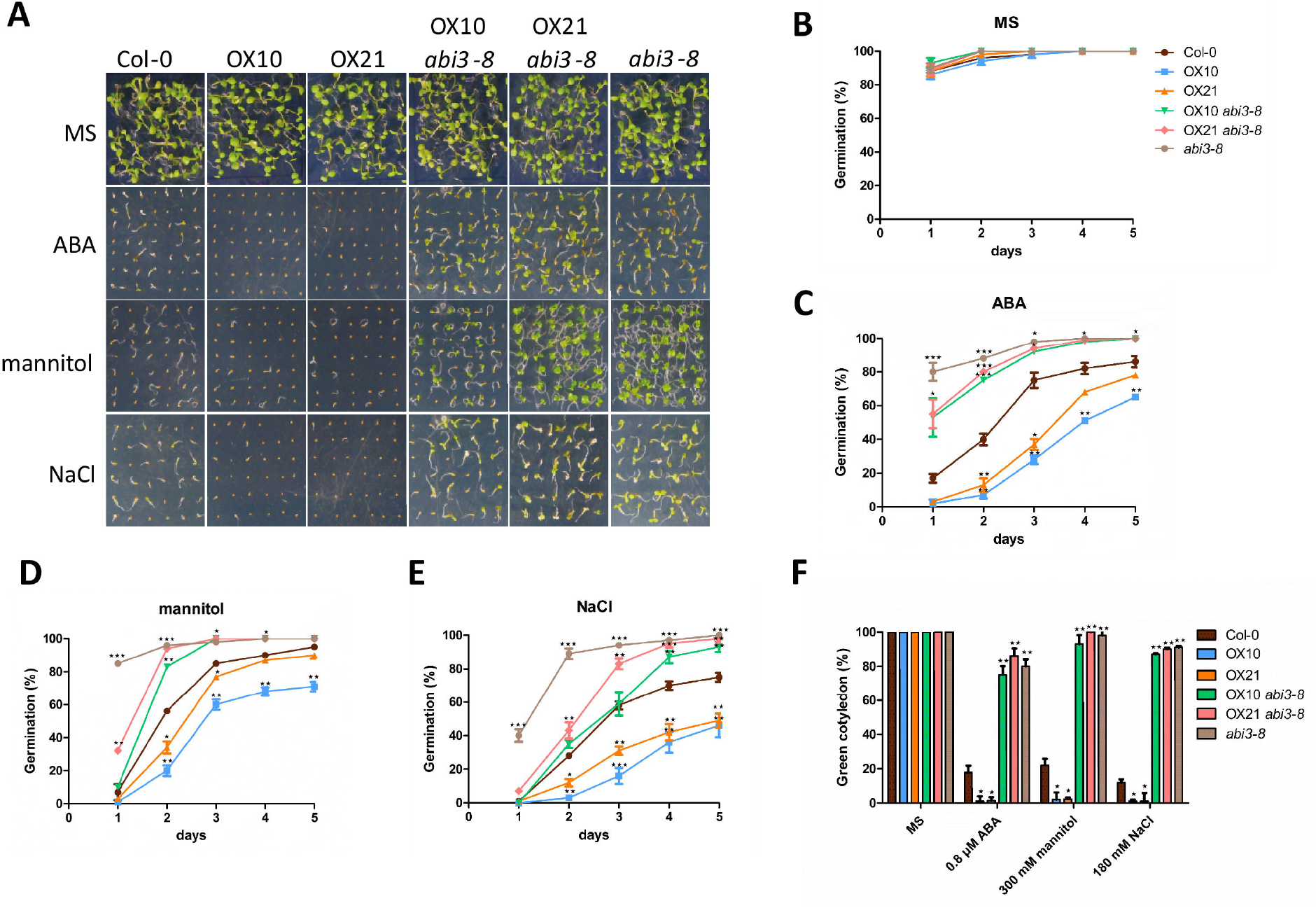
Seed germination response of ANR1-overexpressing lines and *ANR1-OX abi3-8* double mutant to ABA, mannitol and NaCl. A. Phenotypic comparison. Vernalized seeds were transferred to MS medium or MS medium containing 1.0 μM ABA, 300 mM mannitol and 180 mM NaCl and grown for 8 days before the images were recorded. B-E. Seed germination curves. Seed germination rates were calculated at the indicated time for the lines and treatments in A. At least 36 seeds per genotype were measured in each replicate. Data are shown as mean ± SE (n = 3). F. Cotyledon greening ratio. Green cotyledon ratios were counted at day 12 after the end of seed imbibition. Data are shown as mean ± SE (n = 3). *P < 0.05, **P < 0.01.

### ANR1 positively regulates ABI3 protein accumulation

ABI3 protein content adversely affects ABA-dependent seed germination and early-germination growth. To study whether ANR1 also affect ABI3 protein level in seeds besides the upregulated transcription, we detected ABI3 protein content in the wild type, *ANR1*-overexpressing lines, and *anr1-1* seeds by western blot analyses. There were no significant differences in ABI3 protein level between seeds of different genotypes germinated on MS medium for 2 (Figure 7A and B) and 3 days (Figure 7C and D). However, in the presence of ABA, ABI3 protein levels was significantly elevated in *ANR1*-overexpressing seeds, while significantly dropped in *anr1-1* seeds compared with that of wild type seeds (Figure 7). These results are in line with the transcript data and further suggest that ANR1 regulates seed germination through modulating ABI3.

**Figure 7.**
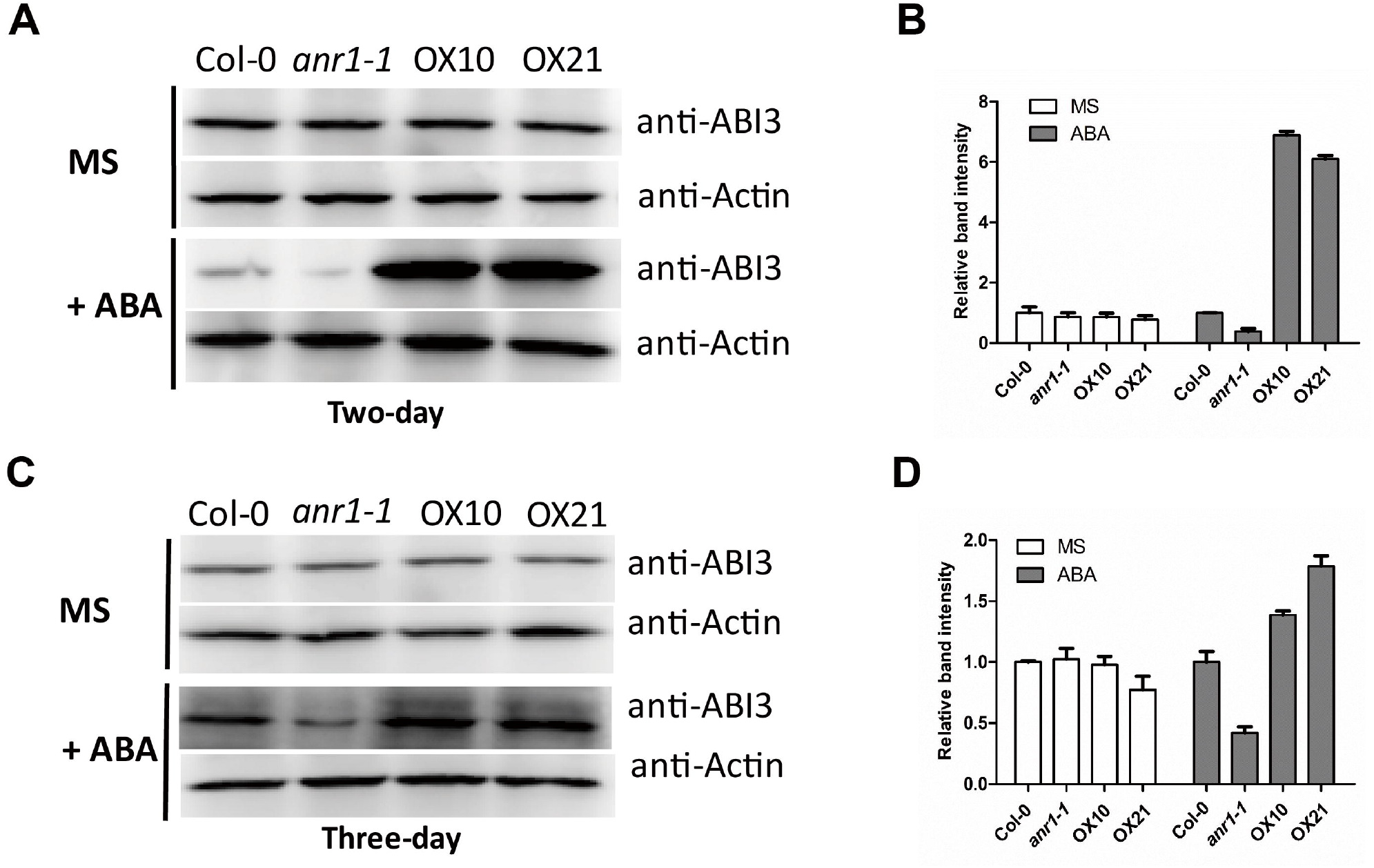
ANR1 positively regulates *ABI3* protein accumulation. A-D. *ABI3* protein levels in wild type (Col-0), *ANR1*-overexpressing, and *anr1-1* germinating seeds. Vernalized seeds (4°C for 3 days) were germinated on MS medium or MS supplement with 0.2 μM ABA for 2 or 3 days. Total protein was extracts and separated by 10% SDS-PAGE and then used for western blot analysis with anti-ABI3 antibody. The Actin protein was used as an internal control. Relative band intensity was measured using ImageJ software (NIH, USA). Data are shown as mean ± SE (n = 3).

### ANR1 up regulates *ABI3* expression by directly binds to its promoter

In plants, MADS-box transcription factors are known to bind the CArG-box (C-[A/T]rich-G) *cis* element within the promoters of their target genes to regulate their expression (Riechmann *et al*., 1996). Through promoter sequence analysis, we found several candidate CArG-box motifs within the *ABI3* promoter (Figure 8A), which led us to examine whether ANR1 could directly bind to the *ABI3* promoter to regulate its expression by chromatin immunoprecipitation (ChIP) assay. The results show that, in the four promoter region checked, only the region containing *cis2* was enriched in ChIP assays with the anti-HA antibody, and the enrichment was enhanced with ABA treatment (Figure 8B). Moreover, we confirmed this result by yeast-one-hybrid assay. Figure 8C shows that ANR1 protein specifically bound to the ABI3 promoter region containing *cis2* in yeast. All these results demonstrate that ANR1 can directly regulate *ABI3* expression by binding to its promoter.

**Figure 8.**
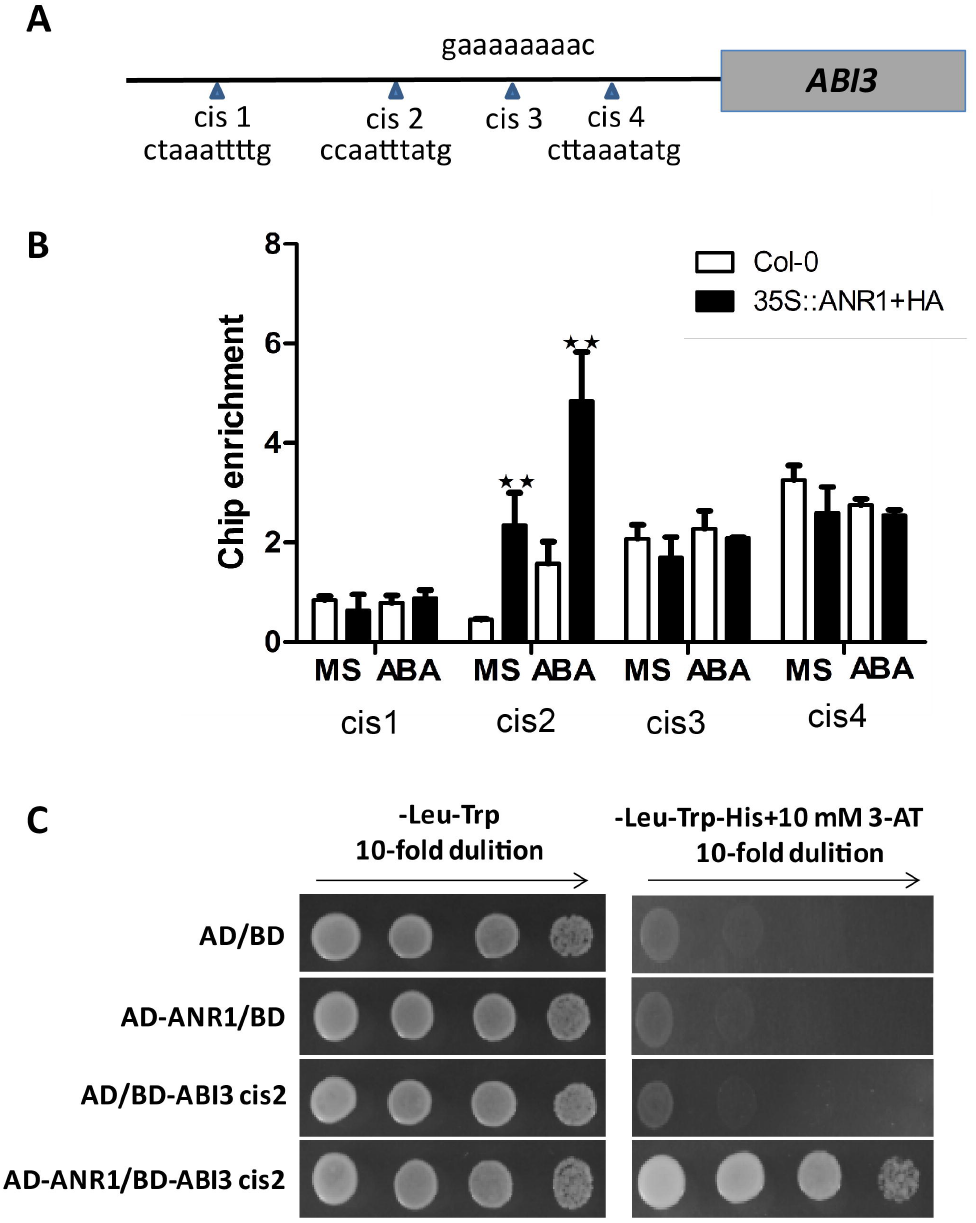
ANR1 directly binds to *ABI3* promoter. A. Illustration of *ABI3* promoter with relative positions of predicted CArG-box motifs, which are indicated with triangles. B. ChIP-qPCR assay of ANR1 binding to *ABI3* promoter. The 7-day-old wild type (Col-0) and 35S:ANR1-HA (OX21) seedlings were transferred to MS solution with or without 20 μM ABA for 6 h and then the chromatins were harvested for ChIP-qPCR assay. Chromatins were immunoprecipitated with anti-HA antibody, and amount of indicated *cis* element in immune complex was detected by qRT-PCR. *UBQ5* was used as an internal reference. Biological triplicates were averaged. Data are shown as mean ± SE (n = 3). *P < 0.05, **P < 0.01. C. Yeast-one-hybrid assay for ANR1 binding to the 30bp sequence of *ABI3* promoter. A serial yeast dilutions (1:1, 1:10, 1:100 and 1:1,000) were grown on SD medium (-Leu-Trp), and the same SD medium minus His (-His-Leu-Trp). The empty pAD and pHIS2 vectors were used as negative control.

### ANR1 and AGL21 function synergistically during seed germination

As the ANR1 and AGL21 belong to the same subfamily with high similarity of amino acid sequence, we wanted to know whether they have functional redundancy in seed germination. We obtain *agl21-1 anr1-1* double mutant by crossing and found that the *agl21-1 anr1-1* seeds showed much higher germination rate and green cotyledon ratio under 1 μM ABA compared with the single mutants, suggesting that these two genes have functional redundancy and act synergistically to regulate seed germination (Figure 9).

**Figure 9.**
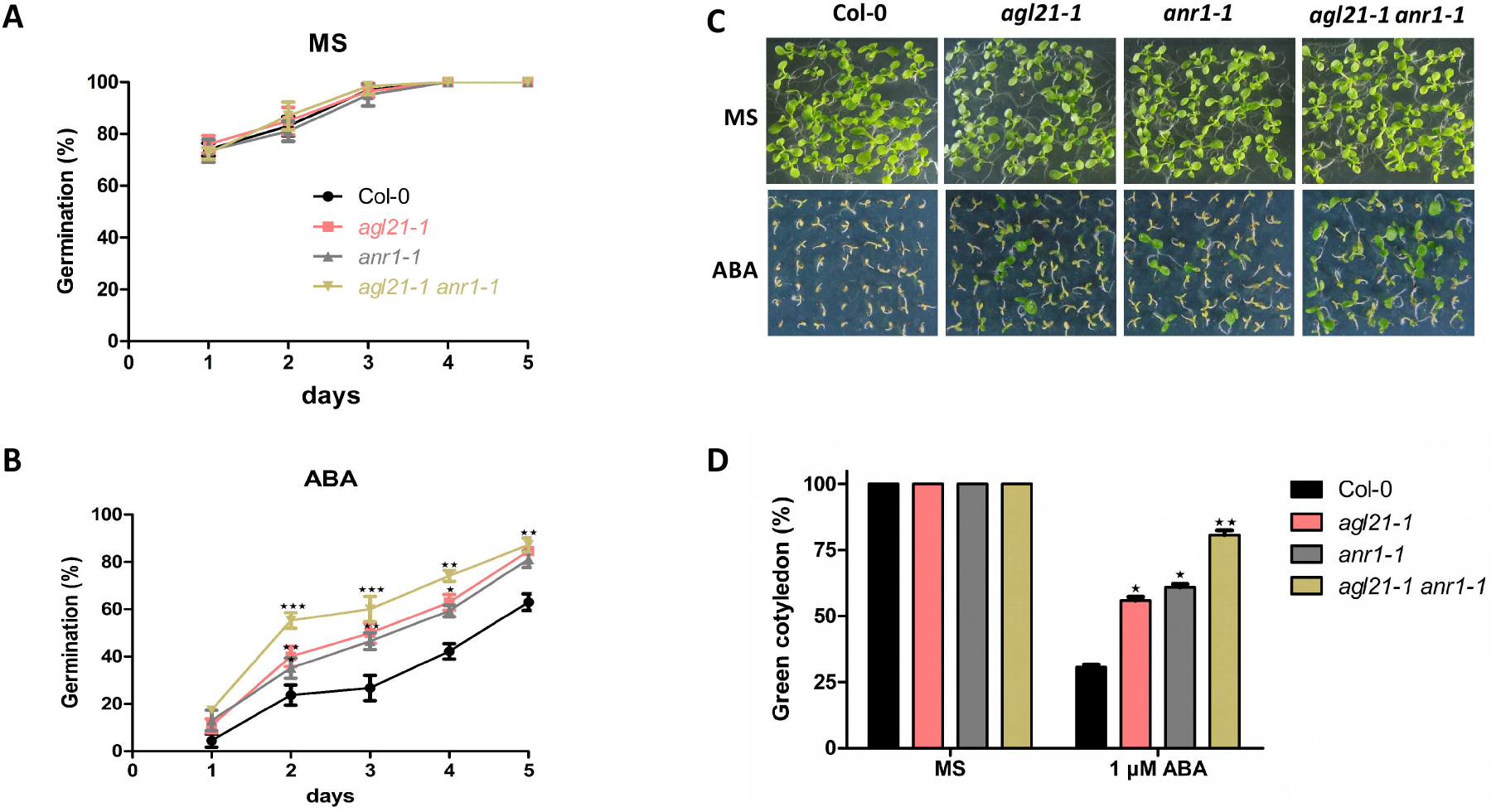
Response of *agl21-1 anr1-1* double mutant seeds to ABA in seed germination. A-B. Seed germination assay. Vernalized seeds (4°C for 3 days) of each genotype were incubated on MS medium or MS supplement with 1.0 μM ABA for 5 days. Seed germination rates were calculated at the indicated time. At least 56 seeds per genotype were used in each replicate. Data are shown as mean ± SE (n = 3). *P < 0.05, **P < 0.01, ***P < 0.001 C. Phenotypic comparison. Vernalized seeds of wild type (Col-0), *agl21-1, anr1-1*, and *agl21-1 anr1-1* double mutant were transferred to MS medium or MS medium containing 1.0 μM ABA, and grown for 12 days before the images were recorded. D. Cotyledon greening ratio. Green cotyledon ratios were counted at day 12 after the end of seed imbibition. Data are shown as mean ± SE (n = 3). *P < 0.05, **P < 0.01.

## DISCUSSTION

ABA is an important stress response hormone, which regulates many important aspects of plant development, including the synthesis of seed storage proteins, promotion of seed desiccation tolerance, and the inhibition of the phase transitions from embryonic to germinative growth (Leung & Giraudat, 1998; Rock, 2000; Rohde *et al*., 2000; Cutler *et al*., 2010). A large number of transcription factors, such as *ABI3, ABI4*, and *ABI5, ELONGATED HYPOCOTYL 5* (*HY5*), *MORE AXILLARY BRANCHES 2* (*MAX2*), *FY, MYB96, WRKY2, WRKY6* have been shown to mediate seed germination by regulating ABA signaling (Parcy *et al*., 1994; Finkelstein *et al*., 1998; Lopez-Molina *et al*., 2002; Chen & Xion, 2008; Jiang & Yu, 2009; Jiang *et al*., 2012; Bu *et al*., 2014; Lee *et al*., 2015; Huang *et al*., 2016). As an important superfamily of transcription factor, MADS-box genes are widely present in eukaryotes and has been extensively explored in diverse developmental processes in flowering plants ranging from root to flower and fruit development (Gramzow & Theissen, 2010). However, their roles in ABA signaling and seed germination remains largely unknown.

In this study, we have demonstrated that MADS-box transcription factor ANR1, which is required for root plasticity in response to nitrate (Zhang and Forde, 1998), is involved in ABA-mediated seed germination and early seedling growth. *ANR1* is highly expressed in dry seeds besides in roots, and it dropped gradually to very low levels during imbibition and germination. Moreover, *ANR1* expression is significantly induced by phytohormone ABA, IAA and MeJA, as well as several environmental stresses such as salt, osmotic, and nitrogen starvation (Figure 1). These data indicate a potential role of ANR1 in seeds germination. Further germination assays showed that ANR1 acts as a negative regulator of seed germination in response to ABA and stresses, as the *ANR1* knockout mutants are insensitive, whereas *ANR1*-overexpression lines are hypersensitive to ABA, salt and osmotic stresses during seed germination and post-germination growth (Figure 2). Based on our data of expression levels of ABA and GA biosynthesis and catabolism pathway-related genes during seed germination and content of ABA and GA in dry and imbibed seeds (Supplemental Figure S2 to S7), ANR1-regulated seed germination may not through affecting ABA or GA content in seeds. However, ANR1 positively modulates multiple ABA-responsive genes, including *ABI3* and *ABI5* (Figure 4), implying that ANR1 may be involved in ABA signaling pathway. We further analyzed the expression levels of ANR1 in germinating *abi* mutant seeds, and found that ANR1 expression could be induced by ABA in *abi3, abi4* and *abi5* mutants, but not in *abi1* and *abi2* mutants (Figure 3). In addition, ANR1 does not affect the expression of upstream ABA signaling pathway genes such as SnRKs, ABI1 and ABI2 (Figure 4). Taken together, these data indicate ANR1 play a role in ABA signaling pathway, acting downstream of SnRKs and PP2Cs but upstream or in parallel with ABI3 and ABI5.

The transcription factors ABI3, ABI4, and ABI5 are well known positive regulators of ABA signaling during seed germination (Giraudat *et al*., 1992; Finkelstein *et al*., 1998; Finkelstein & Lynch, 2000). Previous study reported that ABI3 is an ABA-induced seed dormancy sensor and one of the main regulators of seed embryo maturation as well as early seedling development (Nambara *et al*., 1995). As a key player in ABA-triggered arrest of germination and post germination growth, ABI3 is regulated at both transcriptional and post-transcriptional levels. Several genes, such as *HISTONE DEACETYLASE 6/HISTONE DEACETYLASE 9* (*HDA6/HDA19*), *SUPPRESSOR OF ABI3-5* (*SUA*), *RELATED TO ABI3/VP11* (*RAV1*), *BEL1-LIKE HOMEODOMAIN1*(*BLH1*), *KNOTTED-LIKE FROM ARABIDOPSIS THALIANA 3* (*KNAT3*), *AUXIN RESPONSE FACTOR10/AUXIN RESPONSE FACTOR16* (*ARF10/ARF16*), were reported to regulate seed germination by modulating *ABI3* expression directly or indirectly (Tanaka *et al*.,2008; Sugliani *et al*., 2010; Kim *et al*., 2013; Liu *et al*., 2013; Feng *et al*., 2014). Based on our data, *ABI3* is a potential target of ANR1.

Firstly, the qRT-PCR results show that *ABI3* transcript levels in dry and imbibed seeds are significantly up-regulated in *ANR1*-overexpressing lines while dramatically down-regulated in *anr1* mutants (Figure 4E and 5A). Secondly, we found that *ANR1* and *ABI3* had similar expression patterns in imbibed and germinating seeds (Figure 5B and 5C). Thirdly, we found that loss-of-ABI3 abolished the ABA hypersensitive phenotype of *ANR1*-overexpressing lines in the double mutant *ANR1-OX abi3-8* (Figure 6), indicating that ANR1 regulates seed germination mainly through the regulation of *ABI3*. Moreover, western blot analysis revealed that ANR1 positively modulates ABI3 protein content in germinating seeds in response to exogenous ABA treatment (Figure 7), consistent with the transcription data. To verify whether ANR1 can directly regulate *ABI3* expression, we carried out ChIP and Y1H analysis, and the results show that ANR1 can directly bind to the *ABI3* promoter region containing CArG-box *cis* element (Figures 8). In summary, these data demonstrate that ANR1 is a positive regulator of ABA signaling and suppresses seed germination by directly modulating *ABI3* expression.

Dormancy breaking and germination are vulnerable stages in plant life cycle which are affected by the environmental conditions. Environmental factors such as temperature, light, water, salinity, soil pH and nutrient have great impacts on seed germination. These environmental signals can be perceived by seeds and integrated to endogenous phytohormone signaling, especially ABA and GA (Shu et al., 2016). A number of genes have been found to respond to the environmental stresses and regulate seed germination by integrating the environmental signals to ABA signaling (Vishal & Kumar, 2018). In this study, we found that *ANR1*, like *AGL21* (Yu et al., 2017), also responds to multiple environmental signals, and may act as another environmental surveillance for seed germination. As two members of the AGL17-clade MADS-box genes, *ANR1* and *AGL21* play similar functions during root development as well as seed germination. Together they act as double insurance to modulate plant development to cope with adverse environmental conditions.

In conclusion, we have revealed a novel role of ANR1 in seed germination. ANR1 integrates multiple environmental signals and developmental information to determine the proper timing for seed germination by directly regulating *ABI3* expression. The ANR1-ABI3 signaling module, together with previously reported AGL21-ABI5 module (Yu et al., 2017), forms a safeguard mechanism that double ensures seeds to germinate at right time and under favorable conditions.

## METHODS

### Plant materials and growth conditions

Arabidopsis (*Arabidopsis* thaliana) Columbia-0 ecotype was used for all experiments described. The ANR1 T-DNA insertion lines Salk_052716 and Salk_043618, were ordered from the ABRC. To produce *ANR1*-overexpressing lines (*35S:ANR1-4, 35S:ANR1-10* and *35S:ANR1-21*), the CDS (coding sequence) fragment fused with HA-tag was amplified by PCR and then cloned into the vector pCB2004 through Gateway cloning system. The primer sequences used are listed in Supplemental Table S1. The *ANR1-OX abi3-8* double mutants were generated by genetic cross of *ANR1-OX* and *abi3-8* mutants.

Plants were grown at 22°C under long-day conditions (16-h-light/8-h-dark cycle), and seeds were collected from independent plants at the same time. Harvested seeds were dried at room temperature at least 1 month before germination assays. For the seed germination assay, seeds were surface-sterilized for 15 min in 10% bleach, washed at least four times with sterile water, stratified at 4°C for 48h, and plated on MS solid medium [with 1% (w/v) sucrose] or MS medium supplemented with ABA, NaCl, or mannitol at 22°C under 16-h light/8-h dark photoperiod. For each germination assay, biological triplicates were performed. Seeds from independent plants of the same genotype were used for each replicate. Germination was defined as an obvious emergence of the radicle through the seed coat.

### qRT-PCR analysis

For qRT-PCR analysis, total RNA of seeds was extracted with Trizol reagent (Invitrogen) and RNAs from dry or imbibed seeds at various times. DNase I-treated total RNA (2 μg), and 1 μg of total RNA was used for reverse transcription with oligo(dT)_18_ to synthesize first-strand complementary DNA (cDNA). Real-time qPCR was performed in 48-well blocks with a StepOne Plus Real Time PCR System by using TaKaRa SYBR Premix Ex Taq II reagent kit (TaKaRa, Tokyo, Japan). All qRT-PCRs were performed in triplicate using total RNA samples extracted from three independent biological replicates. The thermal treatment was 5 min at 95°C, then 40 cycles of 10s at 95°C, 20s at 60°C. *UBQ5* was used as an internal control. The primers used are listed in Supplemental Table S1.

### Quantification of ABA and GA

Total free ABA content was measured by ELISA as described by (Yu *et al*., 2017). GAs were extracted and bioactive GA_3_ was quantified according to the method described by Yang et al. (Yang *et al*., 2001). The mouse monoclonal antigens and GA_3_ antibodies, and immunoglobulin G (IgG) horseradish peroxidase were purchased from the Phytohormones Research Institute (China Agricultural University, Beijing, China). ELISA was carried out on a 96-well microtitration plate. The wells were coated with 100 μL of coating buffer (2.93 g L^-1^ NaHCO_3_, 1.5 g L^-1^ Na_2_CO_3_, 0.25 μg mL^-1^ GA antigens, pH 9.6). The coated plates were incubated overnight at 4°C and then kept at room temperature for 30 min. After washing five times with PBS (8 g L^-1^ NaCl, 0.2 g L^-1^ KH_2_PO_4_, 2.96 g L^-1^ Na_2_HPO_4_·12H_2_O, pH7.5) + Tween 20 [0.1% (v/v)] buffer, each well was filled with 50 μL of either GAs standards (0–500 ng mL^-1^ dilution range) or seed extracts, and 50 μL of 20 μg mL^-1^ GAs antibodies, respectively. The plate was incubated at 37°C for 1 h and then washed as described above. One hundred microliters of 1.25 μg mL^-1^ IgG–horseradish peroxidase substrate was added to each well and incubated at 37°C for 0.5 h. The wells were washed five times with the PBS + Tween 20 buffer, and each well was added with 100 μL of color-appearing solution (1.5 mg mL^-1^ o-phenylenediamine, 30% H_2_O_2_). When the 0 ng mL^-1^ standard had a deep color and 500 ng mL^-1^ standard had a pale color in the wells, each well was filled with 50 μL of 2 M H_2_SO_4_ to stop the reaction. Color development in the wells was detected by an ELISA Reader (model EL310, Bio-TEK, Winooski, VT, USA) at optical density A_490_. GA contents were calculated following Weiler et al. (Weiler *et al*., 1981).

### Western blot

To detect the *ABI3* protein levels in ANR1-overexpressing transgenic plants and *anr1* mutants, western blotting was performed according to previously described protocols (Yu *et al*., 2017). Seeds grown on MS medium without or with 0.2 μM ABA for two days and three days were ground in liquid nitrogen and proteins were extracted with RIPA buffer. Protein extracts were separated by 10% SDS-PAGE and transferred onto nitrocellulose membranes. ABI3 protein levels were detected with anti-ABI3 antibody (Abiocode, Agoura Hills, CA, USA). Actin protein levels were used as a control using anti-Actin antibody (Abiocode, Agoura Hills, CA, USA).

### ChIP-qPCR assay

For the ChIP assay, 3 g of 7-day-old 35S:ANR1-HA (OX21) seedlings grown on MS medium was transferred to MS solution with or without 20 μM ABA for 6 h. The ChIP experiment was performed as described previously (Cai *et al*., 2014). Anti-HA antibodies (Abmart, Shanghai, China), and salmon sperm DNA/protein A agarose beads (Millipore, Billerica, MA, USA) were used for ChIP experiments. The enrichment of DNA fragments was analyzed by qPCR using the primers listed in Supplemental Table S1. The purified DNA and input DNA were used as templates. Values were normalized with the level of input DNA. An *UBQ5* fragment was amplified as control.

### Yeast-one-hybrid assay

To test the interaction between ANR1 and the corresponding cis-element of *ABI3*. We used pAD-GAL4-2.1 and the reporter plasmid pHIS2. To obtain pAD/ANR1 plasmid, the CDS (coding sequence) fragment of ANR1 was amplified by PCR and then cloned into the vector pAD-GAL4-2.1 (AD vector) using BamHI and Sall sites. Then, the *ABI3* cis2 element were synthesized with cohesive ends. This sequence was annealed and ligated into the Sacl and MIul sites of the reporter plasmid pHIS2 (BD vector). Yeast cells were grown on two-deficiency (-Trp-Leu) medium for 3 days at 30°C. Then, yeast cells without or with different dilutions (1:10, 1:100 and 1:1,000) were transferred to two-deficiency (-Trp-Leu) medium and three-deficiency (-Trp-Leu-His) medium with 10 mM 3-aminotriazole (3-AT). The pAD and pHIS2 empty vector were used as negative control. The primers we used are listed in Supplemental Table S1.

### Statistical analyses

Statistical analyses were based on Student’s *t*- tests. Values are the mean ± SE (*P <0.05, **P < 0.01, and ***P < 0.001).

### Accession numbers

ANR1: AT2G14210, AGL21: AT4G37940, ABI1: AT4G26080, ABI2: AT5G57050, ABI3: AT3G24650, ABI4: AT2G40220, ABI5: AT2G36270, SnRK2.2: AT3G50500, SnRK2.3: AT5G66880, AtEM1: AT3G51810, RD29A: AT5G52310, RD29B: AT5G52300, ACBP1: AT5G53470, MYB96: AT5G62470, ABA1: AT5G67030, ABA2: AT1G52340, ABA3: AT1G16540, AAO3: AT2G27150, NCED3: AT3G14440. CYP707A1: AT4G19230, CYP707A2: AT2G29090, CYP707A3: AT5G45340, CYP707A4: AT3G19270, CPS: AT4G02780, KS: AT1G79460, KAO2: AT2G32440, GA20ox1: AT4G25420, GA20ox2: AT5G51810, GA20ox3: AT5G07200, GA3ox1: AT1G15550, GA3ox2: AT1G80340, GA3ox3: AT4G21690, GA2ox1: AT1G78440, GA2ox2: AT1G30040, GA2ox3: AT2G34500, GA2ox4: AT1G47990, GA2ox6: AT1G02400, GA2ox7: AT1G56960, GA2ox8: AT4G21200, GAMT1: AT4G26420, GAMT2: AT5G56300, GID1a: AT3G05120, GID1b: AT3G63010, GID1c: AT5G27320, GAI: AT1G14920, RGA: AT2G01570, RGL1: AT1G66350, SPY: AT3G11540, SLY1: AT4G24210, SNE: AT5G48170.

## SUPPLEMENTAL INFORMATION

Supplemental Figure S1. Expression analyses of *ANR1*-overexpressing and *anr1* mutant plants.

Supplemental Figure S2. Expression Levels of ABA biosynthesis and catabolism genes in *anr1* mutants and *ANR1*-overexpressing lines.

Supplemental Figure S3. Response of *anr1* mutants and *ANR1*-overexpressing lines to GA in seed germination.

Supplemental Figure S4. Expression levels of GA synthesis genes in *anr1* mutants and *ANR1*-overexpressing lines.

Supplemental Figure S5. Expression levels of GA catabolism pathway genes in *anr1* mutants and *ANR1*-overexpressing lines.

Supplemental Figure S6. Expression levels of GA signaling pathway genes in *anr1* mutants and *ANR1*-overexpressing lines.

Supplemental Figure S7. ABA content and GA content in WT, *anr1* mutant and anr1-overexpressing plants.

Supplemental Table S1. Primers used for PCR.

## AUTHOR’S CONTRIBUTION

J.L., L.Y. conducted experiments. J.L., L.Y. wrote the manuscript. L.Y. and C. X. edited the manuscript. C. X. supervised the project.

## ACKNOLLEDGEMENT

This work was supported by grants from NNSFC (31572183, 31500231) and MOST (2018ZX08009-11B, 2016ZX08005-004-003, 2016ZX08001003). The authors thank the ABRC for *ANR1* T-DNA insertional lines.

