## supplemental info for "ARABIDOPSIS NITRATE REGULATED 1 acts as a negative modulator of seed germination by activating *ABI3* expression"

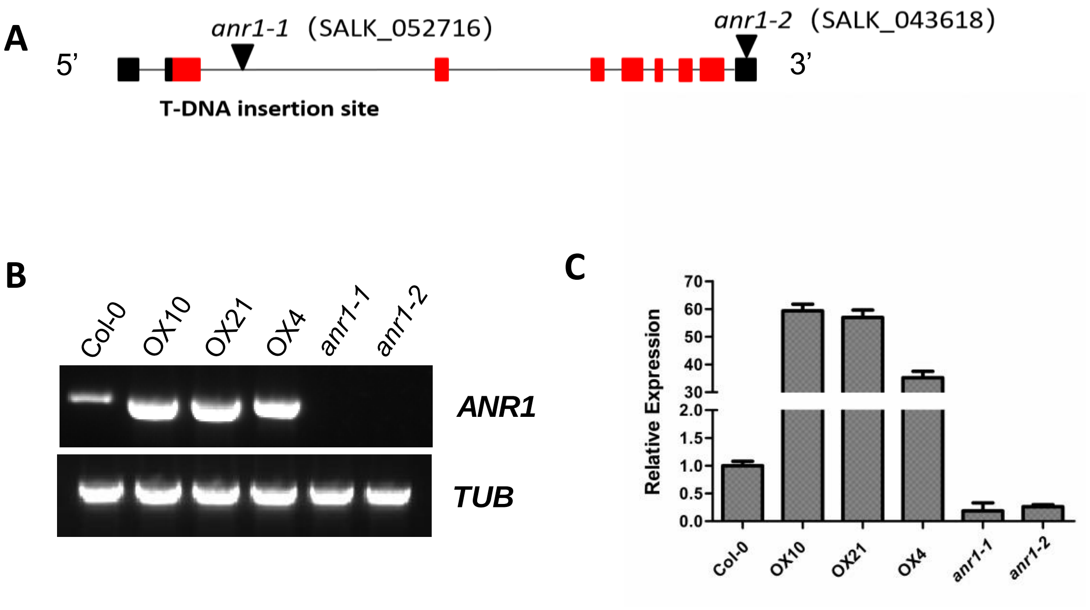

**Supplemental Figure S1. Expression analyses of *ANR1*-overexpressing and *anr1* mutant plants.**

**A**, Mapping of T-DNA insertion sites in *anr1* mutants. Exons are shown as boxes and introns are shown as lines. The triangles indicate the position of the T-DNA insertion.

**B**, Semi-quantitative RT-PCR analyses of *ANR1* expression in Col-0 and *anr1* mutants (*anr1-1*, *anr1-2*) and *ANR1*-overexpressing lines (OX10, OX21, OX4). 5-day-old seedlings were used for RT-PCR analyses using primers *ANR1* CDS LP and *ANR1* CDS RP. *TUBULIN (TUB)* was used as the internal control.

**C**, Expression of *ANR1* was analyzed by qRT-PCR in the *anr1* mutants and *ANR1*-overexpressing lines. Data are shown as mean ± SE (n = 3).

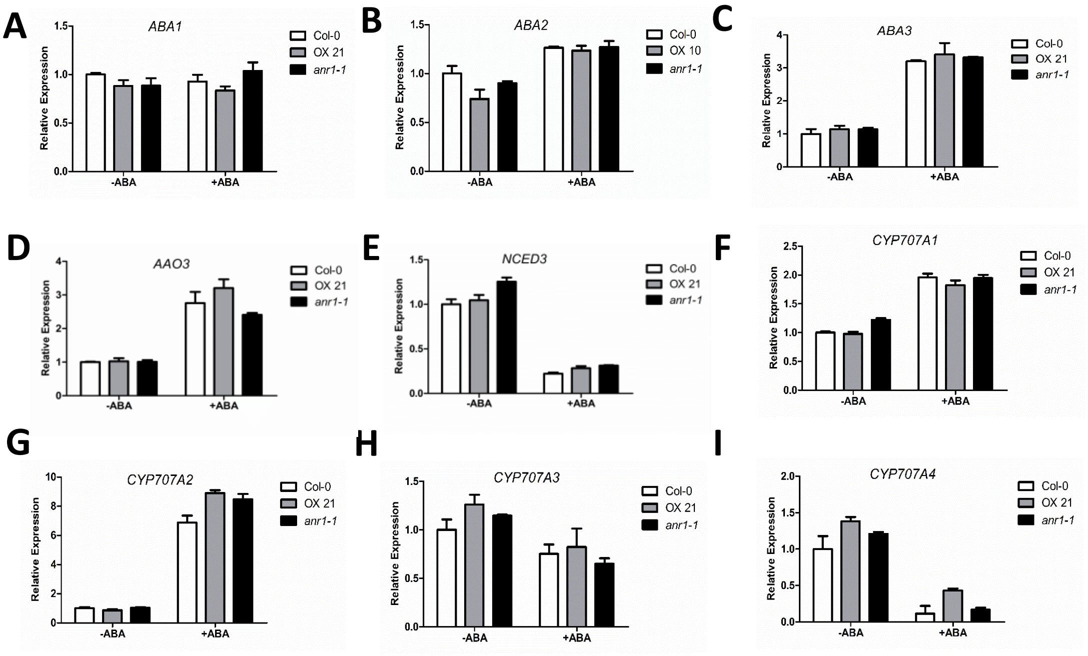

**Supplemental Figure S2. Expression Levels of ABA biosynthesis and catabolism genes in *anr1* mutants and *ANR1*-overexpressing lines.**

The imbibed seeds were germinated and grown on MS medium supplemented with or without 1 μM ABA and incubated for 3 d, and then the plants were harvested for RNA extraction. Transcript accumulation was analyzed by qRT-PCR. The *UBQ 5* gene was used as an internal control. Data are shown as mean ± SE (n = 3). *P < 0.05, **P < 0.01.

A-E, Expression levels of ABA biosynthesis genes in *anr1* mutants and *ANR1*-overexpressing lines.

F-I, Expression levels of ABA catabolism genes in *anr1* mutants and *ANR1*-overexpressing lines.

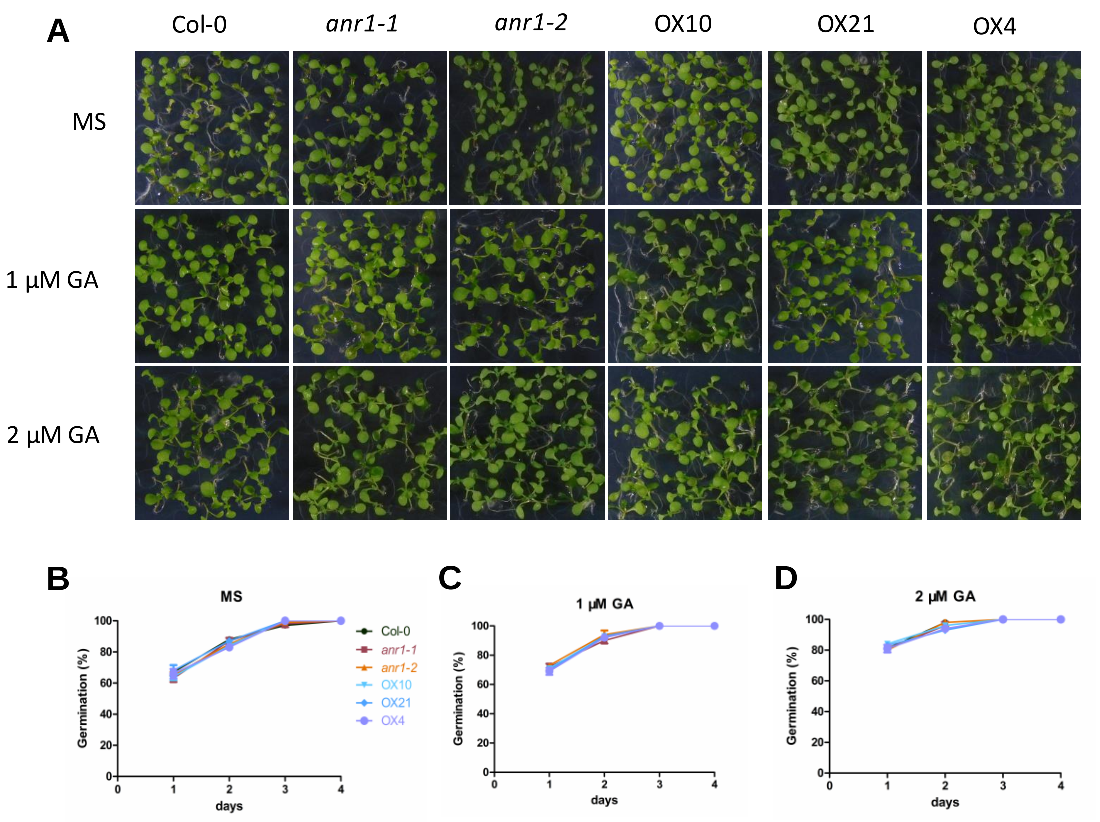

**Supplemental Figure S3. Response of *anr1* mutants and *ANR1*-overexpressing lines to GA in seed germination.**

A, Phenotypic comparison. Imbibed seeds were transferred to MS medium or MS medium containing 1 μM GA, 2 μM GA and grown for 4 d.

B-D, Seed germination assay. Seed germination rates were calculated at the indicated time. At least 42 seeds per genotype were measured in each replicate. Data are shown as mean ± SE (n = 3).

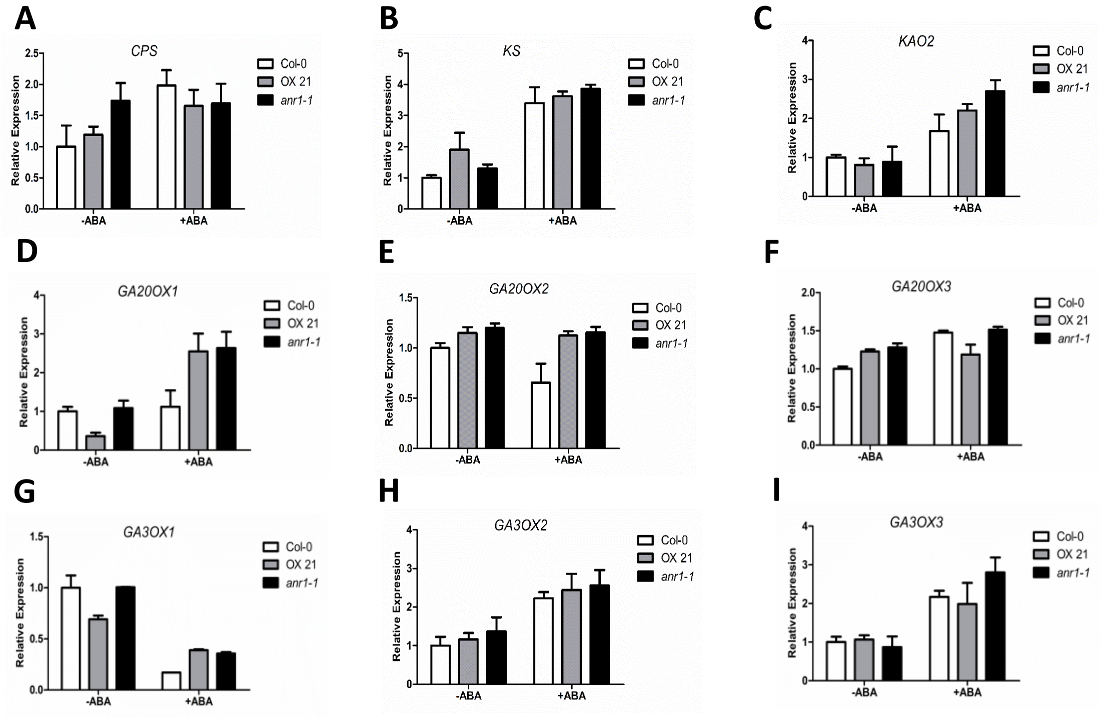

**Supplemental Figure S4. Expression levels of GA synthesis genes in *anr1* mutants and *ANR1*-overexpressing lines.**

A-I, Expression levels of GA synthesis genes. The imbibed seeds were germinated and grown on MS medium supplemented with or without 1 μM ABA and incubated for 3 d, and then the plants were harvested for RNA extraction. Transcript level was analyzed by qRT-PCR. The *UBQ5* gene was used as an internal control. Data are shown as mean ± SE (n = 3). *P < 0.05, **P < 0.01.

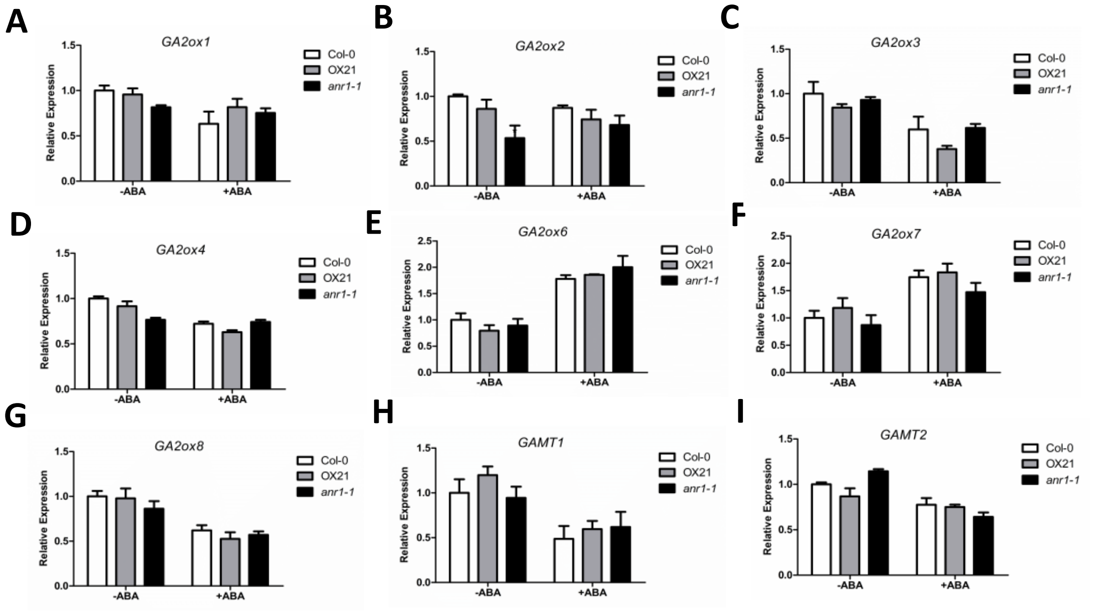

**Supplemental Figure S5. Expression levels of GA catabolism pathway genes in *anr1* mutants and *ANR1*-overexpressing lines.**

**A-I,** Expression levels of GA catabolic pathway genes. The imbibed seeds were germinated and grown on MS medium supplemented with or without 1 μM ABA and incubated for 3 d, and then the plants were harvested for RNA extraction. Transcript accumulation was analyzed by qRT-PCR. The *UBQ 5* gene was used as an internal control. Data are shown as mean ± SE (n = 3).

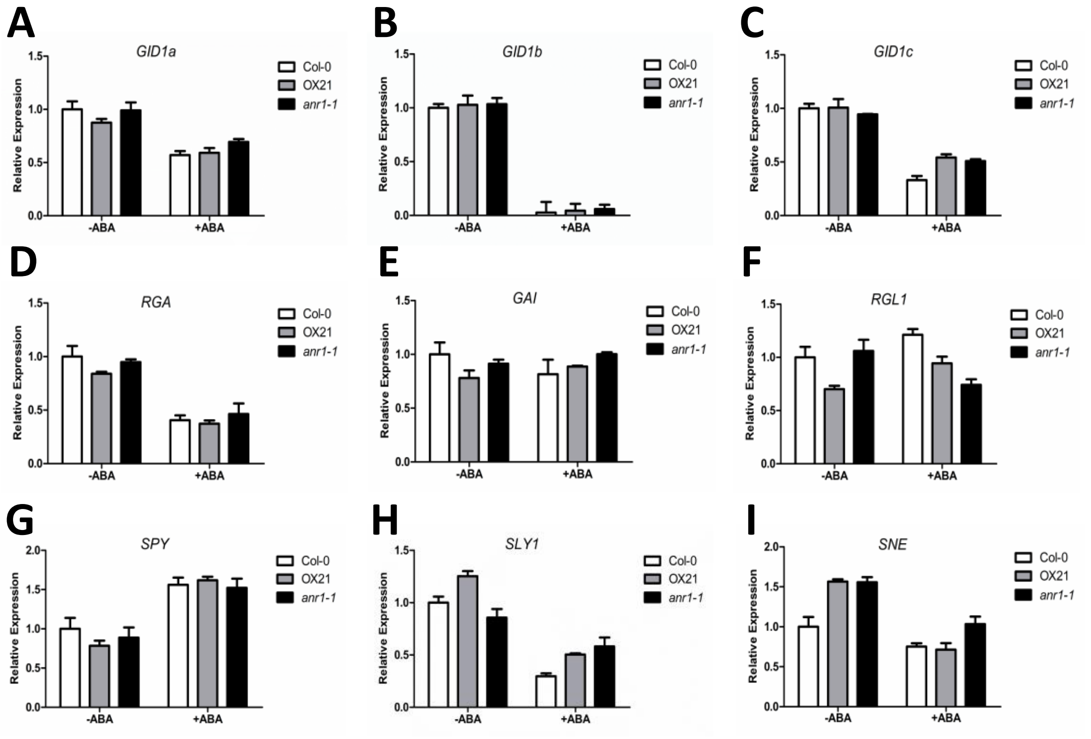

**Supplemental Figure S6. Expression levels of GA signaling pathway genes in *anr1* mutants and *ANR1*-overexpressing lines.**

**A-K,** Expression levels of GA signaling pathway genes. The imbibed seeds were germinated and grown on MS medium supplemented with or without 1 μM ABA and incubated for 3 d, and then the plants were harvested for RNA extraction. Transcript accumulation was analyzed by qRT-PCR. The *UBQ 5* gene was used as an internal control. Data are shown as mean ± SE (n = 3). *P < 0.05.

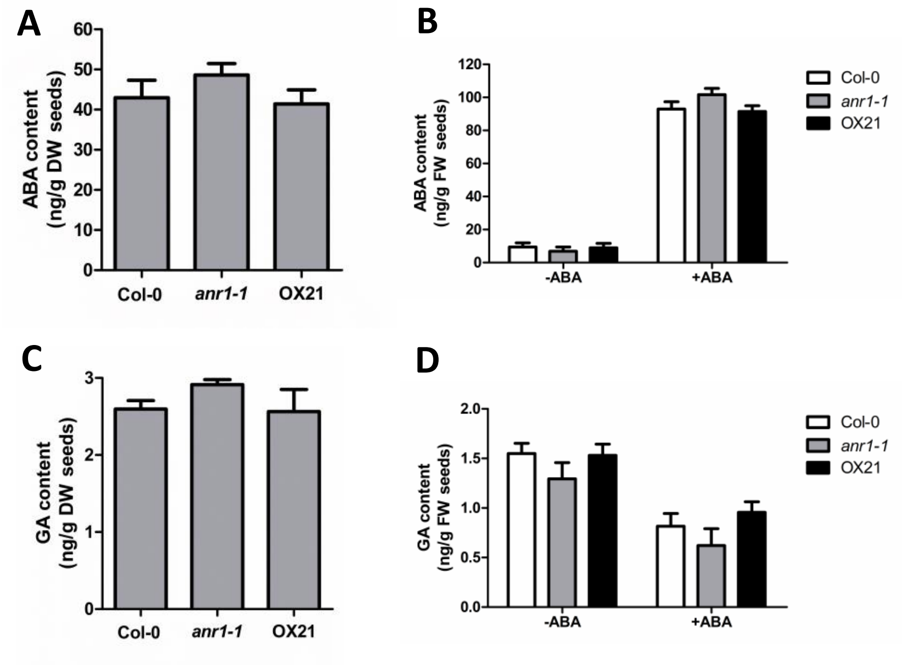

**Supplemental Figure S7. ABA content and GA content in WT, *anr1* mutant and ANR1-overexpressing plants.**

The dry seeds and imbibed seeds were germinated and grown on MS medium supplemented with or without 1 μM ABA and incubated for 3 d, and then the plants were harvested for ELISA.

A and C, ABA and GA contents in dry seeds of the wild type (Col-0), *anr1* mutant and *ANR1*-overexpressing plants. Three independent experiments were conducted. Values are mean ± SD of three replications.

B and D, ABA and GA contents in imbibed seeds of the wild type (Col-0), *anr1* mutant and *ANR1*-overexpressing plants. Three independent experiments were conducted. Values are mean ± SD of three replications.

**Supplemental Table S1. Primers used for PCR**

| **Name** | **Sequence (5' to 3')** |
|---|---|
| *ANR1-HA* LP | GGGGACAAGTTTGTACAAAAAAGCAGG CTATGGGGAGAGGGAAGATAGTT |
| *ANR1-HA* RP | GGGGACCACTTTGTACAAGAAAGCTGGGT CTAGGAAAGTTGTAGCCCTAG |
| *ANR1*-CDS LP | ATGAGGAAATCCAAAGCTAAT |
| *ANR1*-CDS RP | CTAGGAAAGTTGTAGCCCTAG |
| *TUB LP* | CTTAAGCTCACCACTCCAAGCT |
| *TUB RP* | GCACTTCCACTTCGTCTTCTTC |
| Salk_052716 LP | TTCCATTTATTGCAAACCCTG |
| Salk_052716 RP | AATGCGCATGAATGTCTTTTC |
| Salk_043618 RP | ATGGAAGAACAAATGCGATTG |
| Salk_043618 LP | TGTTCAGTTGGGACAATCTCC |
| Primers for qRT-PCR | |
| *ANR1*qPCR LP | CTTCATGCTGGAGCTTGCAAAGTC |
| *ANR1*qPCR RP | AGCTATTCTCTGTGATGCCGAGGT |
| *UBQ5* qPCR LP | AGAAGATCAAGCACAAGCAT |
| *UBQ5* qPCR RP | CAGATCAAGCTTCAACTCCT |
| *ABA1* qPCR LP | GCTACTCTATCAATCCATCTCC |
| *ABA1* qPCR RP | CTCTCTTCTCCTCCTTCTCAAC |
| *ABA2* qPCR LP | GGTGAGAGCATTGTTCGTCTG |
| *ABA2* qPCR RP | ACGGTGCTCCACATAATCCTG |
| *ABA3* qPCR LP | TACCCAGATGGTCCCAAGAAC |
| *ABA3* qPCR RP | CATCCGCTATAAGGTCACTGG |
| *AAO3* qPCR LP | ACGACCTTACTTGAGTTCTTGC |
| *AAO3* qPCR RP | TTCGTGTTTCCAAGACCTTCAG |
| *NCED3* qPCR LP | ATGGCTTCTTTCACGGCAACG |
| *NCED3* qPCR RP | GCGGGAGAGTTTGATGATTGC |
| *CYP707A1* qPCR LP | CCATCGCTCAAGACTCTCTC |
| *CYP707A1* qPCR RP | CCTCGTCTTTTCCGAAGATCG |

| | |
|---|---|
| *CYP707A2* qPCR LP | CAATTCCTTCTTCGCCACTCG |
| *CYP707A2* qPCR RP | GCCTCTGGTCCAATCATACG |
| *AtGA3ox1* (At1g15550) qPCR LP | CCATTCACCTCCCACACTCT |
| *AtGA3ox1* (At1g15550) qPCR RP | GCCAGTGATGGTGAAACCTT |
| *AtGA3ox2* (At1g80340) qPCR LP | TGGTCCGAAGGTTTCAC |
| *AtGA3ox2* (At1g80340) qPCR RP | GGGTCGAGTCTGTATGG |
| *AtGA3ox3* (At4g21690) qPCR LP | TCCTACCCGGTTTGCC |
| *AtGA3ox3* (At4g21690) qPCR RP | ACGGTGCATTGTACTTC |
| *AtCPS*(At4g02780) qPCR LP | CTAAAGCTTACAAGGATACC |
| *AtCPS*(At4g02780) qPCR RP | GACATTGGGAACTCCTCC |
| *AtKS*(At1g79460) qPCR LP | GGATCTTAAATGTGATAGTG |
| *AtKS*(At1g79460) qPCR RP | AAGGAACTGCAGCCTCG |
| *AtKAO2*(At2g32440) qPCR LP | AGAGCTTTGAAGGCAAGG |
| *AtKAO2*(At2g32440) qPCR RP | TCTCTTGTTCTTCCTTAGC |
| *AtGA20ox1*(At4g25420) qPCR LP | CTCATGAATACACGAGCC |
| *AtGA20ox1*(At4g25420) qPCR RP | TGATACACCTTCCCAAATG |
| *AtGA20ox2*(At5g51810) qPCRLP | ATGCTCACCGTTTGATGG |
| *AtGA20ox2*(At5g51810) qPCR RP | CCTTCCCAAACTGCTCG |
| *AtGA20ox3*(At5g07200) qPCR LP | CCTATCTGCATATGGACTC |
| *AtGA20ox3*(At5g07200) qPCR RP | AAACCTTCCCGAAATCTTC |
| *ABI1* qPCR LP | CGGCAAAACTGCACTTCCATT |
| *ABI1* qPCR RP | CACGAGCTCCATTCCACTGAA |
| *ABI2* qPCR LP | CTCGCAATGTCAAGATCCATTGGC |
| *ABI2* qPCR RP | TTACTCGCCGCACTGAAGTCAC |
| *ABI3* qPCR LP | GATTGAATCAGCGGCAAGAA |
| *ABI3* qPCR RP | GTTGTTGTGGTGGTGGAGGA |
| *ABI4* qPCR LP | CTCGCAATGTCAAGATCCATTGGC |
| *ABI4* qPCR RP | TTACTCGCCGCACTGAAGTCAC |
| *ABI5* qPCR LP | CAGCTGCAGGTTCACATTCTG |
| *ABI5* qPCR RP | CACCCTCGCCTCCATTGTTAT |
| *AtEM1* qPCR LP | GGGACGTAAAGGAGGACTCAGTA |
| *AtEM1* qPCR RP | CTTTGACTCATCGATCTCAATCC |
| *RD29A* qPCR LP | CCTGAAGTGATCGATGCACCAG |
| *RD29A* qPCR RP | TGGTGTAATCGGAAGACACGAC |
| *RD29B* qPCR LP | GTGAAGATGACTATCTCGGTGG |
| *RD29B* qPCR RP | CACCACTGAGATAATCCGATCC |
| *SnRK2.2* qPCR LP | TTGTTAGATGGAAGTCCGGCAC |
| *SnRK2.2* qPCR RP | TTGGGAATGAAGAACAGAAGACT |
| *SnRK2.3* qPCR LP | CATACATCGCTCCAGAGGTACT |
| *SnRK2.3* qPCR RP | GATACGCTCCAACCAACATGAC |
| *MYB96* qPCR LP | TGCAGTCTCGGAAGAAGGTG |
| *MYB96* qPCR RP | CATCTCGTGGCTTTGCTCAT |

| | |
|---|---|
| *ACBP1* qPCR LP | AACCACACGACTCAATCGGA |
| *ACBP1* qPCR RP | TTTCGACACCTTCCCAATCA |
| Primers for ChIP-qPCR | |
| *ABI3* ChIP cis1 LP | GTAAAGCAAATAATCCCATAAC |
| *ABI3* ChIP cis1 RP | TGTGAACTTTGTTTTACTATGC |
| *ABI3* ChIP cis2 LP | AACAATGTCATTTAGAATTGGG |
| *ABI3* ChIP cis2 RP | CATGCACGAACATCATCAATA |
| *ABI3* ChIP cis3 LP | TCATTTTGAAGGCAAGGTCGA |
| *ABI3* ChIP cis3 RP | AACATATTGTAGAACAATGTATG |
| *ABI3* ChIP cis4 LP | CTCCTGTACCAACCTTTAGATA |
| *ABI3* ChIP cis4 RP | AACAAAGTTCCTCGAGAGAAAG |
| Primers for Yeast one-hybrid assay | |
| Y1H AD/ANR1 LP | CGGGATCCATGGGGAGAGGGAAGATAGTT |
| Y1H AD/ANR1 RP | ACGCGTCGACCTAGGAAAGTTGTAGCCCTAG |
| Y1H pHIS2/*ABI3* cis2 LP | CAATTTTAATTCCAATTTATGTATGTGTATAA |
| Y1H pHIS2/*ABI3* cis2 RP | CGCGTTATACACATACATAAATTGGAATTAAAATTGAGCT |
